# Dual inhibition of anti-apoptotic proteins BCL-XL and MCL-1 enhances cytotoxicity of Nasopharyngeal carcinoma cells

**DOI:** 10.1101/2021.07.07.451410

**Authors:** Siti Fairus Abdul Rahman, Azali Azlan, Kwok-Wai Lo, Ghows Azzam, Nethia Mohana-Kumaran

## Abstract

One of the many strategies that cancer cells use to evade cell death is through upregulation of the BCL-2 anti-apoptotic proteins. Hence, these proteins have become attractive therapeutic targets. Given that different cell population rely on different anti-apoptotic proteins for survival, it is crucial to determine which proteins are important for NPC survival. Here we determined the survival requirements for the NPC cells using combination of CRISPR/Cas9 technique and pharmacological approaches. A human apoptosis RT^2^ Profiler PCR Array was first employed to profile the anti-apoptotic gene expressions in NPC cell lines HK-1 and C666-1. The HK-1 cells expressed all the anti-apoptotic genes (*MCL-1, BFL-1, BCL-2, BCL-XL, and BCL-w*). Similarly, the C666-1 cells expressed all the anti-apoptotic proteins except *BFL-1* (undetectable level). Notably, both cell lines highly expressed *MCL-1*. Deletion of *MCL-1* sensitized cells to A-1331852 suggesting that MCL-1 and BCL-XL may be important for NPC cell survival. Co-inhibition of MCL-1 and BCL-2 with MCL-1 selective inhibitor S63845 and BCL-2 selective inhibitor ABT-199 inhibited NPC cell proliferation but the effect on cell viability was more profound with co-inhibition of MCL-1 and BCL-XL with S63845 and A-1331852, implying that MCL-1 and BCL-XL are crucial for NPC cell survival. Furthermore, co-inhibition of MCL-1 and BCL-XL inhibited the growth and invasion of NPC spheroids. Deletion of *BFL-1* sensitized cells to A-1331852 suggesting that BFL-1 may play a role in NPC cell survival. Taken together co-inhibition of BCL-XL and MCL-1/BFL-1 could be potential treatment strategies for NPC.

## Introduction

Nasopharyngeal carcinoma (NPC) is rare worldwide but endemic in Asia especially in South-East Asia region (GLOBOCAN 2020). Treating patients with metastatic NPC is often a challenge as patients develop resistance to systemic anti-cancer therapies and retreating local recurrence with radiotherapy have many limitations. Hence, improved treatment strategies are needed for better patient outcome.

The BCL-2 family of proteins are critical regulators of the apoptosis pathway, intrinsically mediated via the mitochondria. The members of the family are divided into pro- and anti-apoptotic proteins (e.g. BCL-2, BCL-XL, BCL-w, MCL-1 and BFL-1) [2]. The anti-apoptotic proteins are upregulated in many cancers and hence have become attractive therapeutic targets. However, different cell population depend on different anti-apoptotic protein(s) for survival. Hence, it is crucial to determine which anti-apoptotic proteins do NPC cells rely for survival so that they can be optimally targeted for better treatment outcome. The Dynamic BH3 profiling [3], the CRISPR/Cas9 genome editing technique [4] and BH3 mimetics given their selectivity in inhibiting the anti-apoptotic proteins (an approach that is known as ‘chemical parsing’) [5] are approaches that can be used to parse the individual contributions of the BCL-2 anti-apoptotic proteins for cell survival. Our previous study showed that ABT-263, a small molecule inhibitor which selectively inhibits BCL-2, BCL-XL and BCL-w, had minimal effect on NPC cells, which made it clear that other anti-apoptotic proteins may be important for NPC cell survival [6].

This study employed combination of the CRISPR/Cas9 technique and BH3 mimetics namely the BCL-2 inhibitor ABT-199 [7], BCL-XL inhibitor A-1331852 [8] and MCL-1 inhibitor S63845 [9] to delineate cellular BCL-2 family dependence profiles, of the NPC cells.

## Material and Methods

Detailed materials and methods are provided in the Supplementary material (Supplementary Materials and methods, S. Fig1 & S. Fig 2). The expression of the BCL-2 anti-apoptotic genes in the NPC cell lines were accessed using the custom RT^2^ Human Apoptosis Profiler PCR array (Qiagen, Hilden, Germany). To delineate the contribution of MCL-1 for cell survival, the HK-1 cells were transfected with two independent single-guide RNAs (sgRNAs) targeting different regions of the human *MCL-1* gene. *MCL-1* mutagenesis was verified by conventional Sanger DNA sequencing (First Base, Singapore) and gene expression of *MCL-1* in the parental and *MCL-1* deleted cells were verified by qPCR performed using the iTaq™ Universal SYBR Green Supermix (BioRad, California, USA), according to manufacturer’s protocol. The *MCL-1* manipulated cells were treated with either ABT-199 or A-1331852 to access the sensitivity of the *MCL-1* deleted cells to the BH3 mimetics. To complement the gene editing study, the individual contributions of the anti-apoptotic proteins for NPC cell survival were parsed using BH3 mimetics. Three NPC cell lines HK1, C666-1 and C17 [10] were treated with BH3 mimetics namely ABT-199 (MedChemExpress, NJ, USA), A-1331852 (MedChemExpress, NJ, USA) and S63845 (MedChemExpress, NJ, USA) either singly or as combination. Cell viability was assessed using the SyBr Green I assay. Combination of A-1331852 and S63845 was later tested on spheroids generated from the HK-1 and C17 NPC cell lines. Upon termination of spheroid assays, spheroids were stained with Calcein AM (ThermoFisher Scientific, MA, USA) to access cell viability and Ethdium-homodimer I (ThermoFisher Scientific, MA, USA) to access cell death. In order to access the contribution of BFL-1 for NPC cell survival, the HK-1 cells were transfected with two independent single-guide RNAs targeting different regions of the human *BFL-1* gene. Validation of *BFL-1* mutagenesis was similar to validation of *MCL-1* mutagenesis. The *BFL-1* manipulated cells were treated with either ABT-199 or A-1331852 to access the sensitivity of the *BFL-1* deleted cells to the BH3 mimetics.

## Results

### Expression of the BCL-2 anti-apoptotic proteins in the NPC cell lines

The basal expressions of all anti-apoptotic genes in the NPC cell lines HK-1 and C666-1 were first determined using a human apoptosis real time PCR array. In the HK-1 cells all the anti-apoptotic genes were detected. Most of the anti-apoptotic genes required more than 20 cycles of PCR amplification to detect except for *MCL-1* which was detectable within 18 cycles of PCR amplification, which indicated high expression level of this gene in the cells (S. Fig 3a). In the C666-1 cells, all of the anti-apoptotic genes were detectable except *BFL-1*. Similar to the HK-1 cells, MCL-1 was detected within 20 cycles (S. Fig 3b). Taken together, data shows that both NPC cells lines expressed all of the anti-apoptotic genes (except for *BFL-1* in the C666-1 cells) and more notably, both NPC cell lines expressed high levels of *MCL-1*.

### Inhibition of MCL-1, BCL-2 or BCL-XL alone is not sufficient to kill NPC cells

Given that the NPC cell lines expressed high levels of *MCL-1*, the gene was deleted in the HK-1 cells to delineate the role of *MCL-1* for cell survival, using the CRISPR/Cas9 technique. The HK-1 cells were transfected with two independent single-guide RNAs (sgRNAs) targeting different regions of the human *MCL-1* gene (hereafter the sgRNAs will be referred to as sg*MCL-1*#1 and sg*MCL-1*#2). The experimental design is shown in Fig. 1a. *MCL-1* mutagenesis was verified by conventional Sanger DNA sequencing and gene expression of *MCL-1* in the parental and *MCL-1* deleted cells were verified by qPCR. The sg*MCL-1*#2 generated higher number of InDels (insertions and deletions of bases in the DNA) compared to sg*MCL-1*#1 (Fig. 1b). The sg*MCL-1*#1 and sg*MCL-1*#2 resulted in 90% and 99% reduction in *MCL-1* expression, respectively (Fig. 1c).

**Fig. 1:**
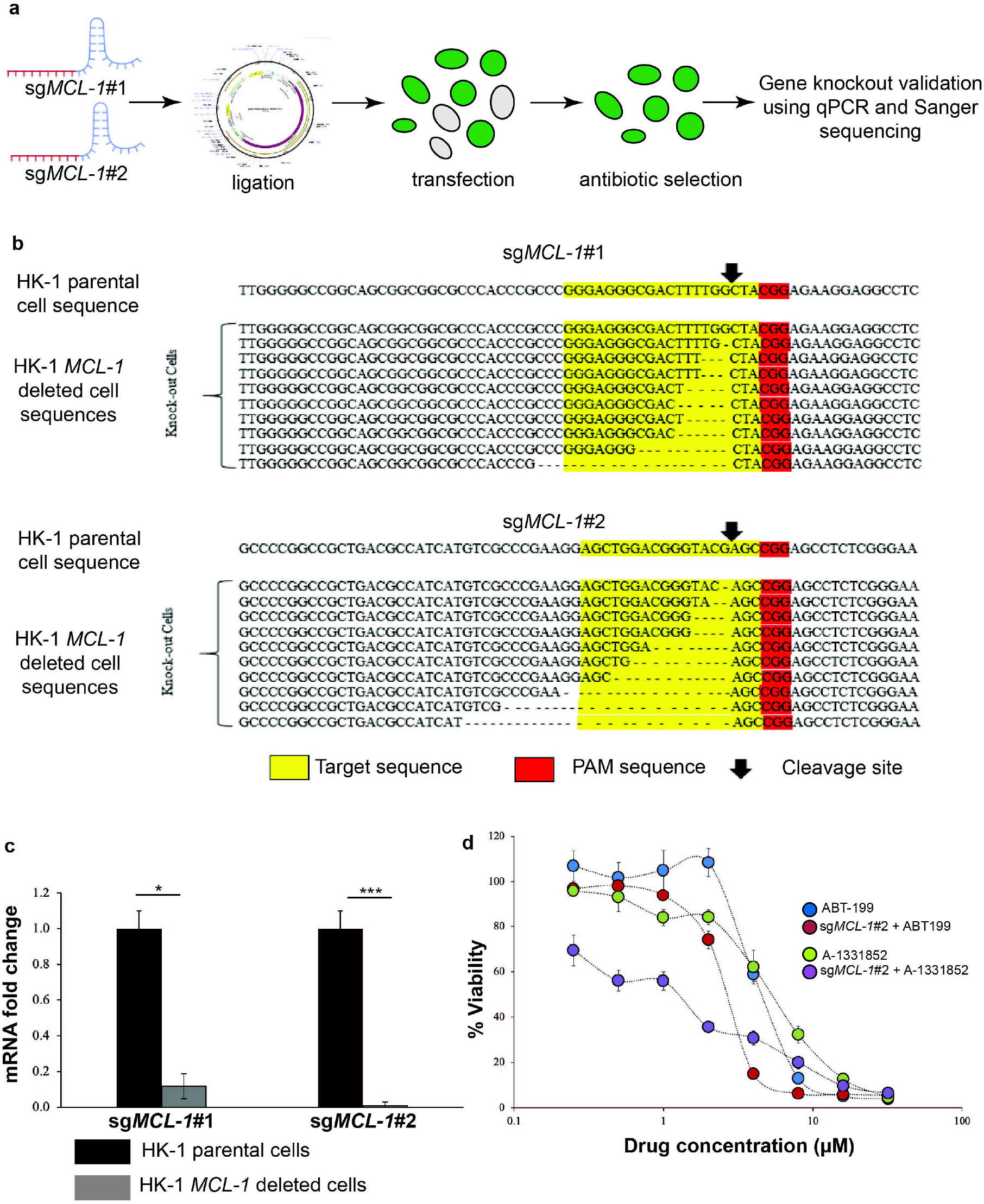
HK-1 *MCL-1* deleted cells were sensitive to treatment of BCL-XL selective inhibitor A-1331852. **(a)** Experimental design to generate the HK-1 *MCL-1* deleted cell lines using the CRISPR/Cas9 technique. **(b)** Read sequences of the HK-1 sg*MCL-1*#1 and HK-1 sg*MCL-1*#2 cells. Parental *MCL-1* sequence is shown on the top. Most of the clones tested showed deletions at the expected cleavage sites (arrow). The dashed lines indicate the InDels. The target sequence is highlighted in yellow while the PAM sequence is highlighted in red. **(c)** qPCR validation *MCL-1* gene deletion in the HK-1 cells. *MCL-1* expression levels were normalized to parental cells. Bars indicate mean SEM of three independent experiments. Statistically significant differences in *MCL-1* expression between the parental cell line and the *MCL-1* knockout cells are shown as ***p ≤ 0.001 or *p < 0.05 determined by two-tailed paired T-test.**(d)** The sensitivity of the HK-1 parental cell line and the HK-1 sg*MCL-1*#2 cells were tested to tested to single agent activity of either ABT-199 or A-1331852 (0-32 µM). Points represent mean SEM of four experiments.

Given that the sg*MCL-1*#2 demonstrated more profound reduction of *MCL-1* expression, the sg*MCL-1*#2 cells were treated with increasing concentrations of either ABT-199 or A1331852 to access whether *MCL-1* deletion sensitized NPC cells to these inhibitors. The HK-1 parental cells were resistant to single agent treatment of ABT-199 and A-1331852 (Fig. 1d). The sg*MCL-1*#2 cells were weakly sensitized to ABT-199 (Fig. 1d – red circle & Table 1). However, the sg*MCL-1*#2 cells were sensitized to A-1331852 by ∼4-fold (Fig. 1d – purple circle & Table 1), indicating that the MCL-1 and BCL-XL may be important for NPC cell survival.

**Table 1:** Sensitivity of the HK-1 NPC cell line to either ABT-199 or A-1331852 following manipulation of *MCL-1*.

| Drug | Cell Type | IC<sub>50</sub> ± SEM (μM) | Fold sensitization |
| --- | --- | --- | --- |
| ABT-199 | Parental HK-1 cell line | 7.11 ± 0.42 |  |
|  | HK-1 sg*MCL-1*#2 cells | 5.18 ± 0.61 | 1.4 |
| A-1331852 | Parental HK-1 cell line | 4.58 ± 0.22 |  |
|  | HK-1 sg*MCL-1*#2 cells | 1.18 ± 0.07 | 3.9 |

NOTE: Values were derived from at least 4 experiments. Sensitization was computed relative to the parent cell line, as shown.

To complement the gene editing study, the individual contributions of MCL-1, BCL-2 and BCL-XL were parsed using BH3 mimetics which selectively inhibit these proteins. The NPC cell lines HK-1 and C666-1 were first treated with single agent ABT-199, A-1331852 and S63845. The BH3 mimetics were also tested on a newly derived NPC cell line, C17. The HK-1 (S. Fig 4a) and C666-1 (S. Fig 4b) cells were resistant to single agent treatment of all three inhibitors. The C17 cells were resistant to single agent treatment of ABT-199 (S. Fig 4c – open square) and A-1331852 (S. Fig 4c – open circle) and but sensitive to single agent treatment of S63845 (S. Fig 4c – open triangle).

Taken together, these data suggest that insensitivity of the HK-1 and C666-1 cells to single agent treatment of all three inhibitors suggest that the cells depend on more than one anti-apoptotic protein for survival. The C17 cells were resistant to single agent treatment of ABT-199 and A-1331852 but the cells were sensitive to S63845 treatment indicating that the cells depend on MCL-1 for survival.

### NPC cell lines were sensitive to co-inhibition of MCL-1 and BCL-2

Given that the NPC cells were insensitive to single agent treatment of the BH3 mimetics, the cells were first tested combination of ABT-199 and S63845. The HK-1 cells were treated with increasing concentrations of ABT-199 (0-32 µM) in the absence and presence of either 0.5, 1 or 2 µM of S63845 for 72 hours. At 0.5 µM S63845, the cells were only sensitized to ABT-199 by 3-fold (S. Fig 5a – open circle & S. Table 1). The sensitization increased to 16-fold at a concentration of 1 µM (S. Fig 5a – open triangle & S. Table 1) and 2 µM of S63845 (S. Fig 5a – open diamond & S. Table 1).

Similarly, the C666-1 cells were treated with increasing concentrations of ABT-199 (0-32 µM) in the absence and presence of either 0.5, 1 or 2 µM of S63845 for 72 hours. At 0.5 µM S63845, the cells were sensitized to ABT-199 by 5-fold (S. Fig 5b – open circle & S. Table 1). The sensitization increased to 12-fold at 1 µM (S. Fig 5b – open triangle & S. Table 1) and increased further to 36-fold at 2 µM of S63845 (S. Fig 5b – open diamond & S. Table 1).

As the C17 cells were sensitive to single agent treatment of S63845 (S. Fig 5c), the sensitivity of the C17 cell lines were tested to fixed concentrations of S63845 (below 1 µM) with increasing concentrations of ABT-199. The C17 cells were treated with increasing concentrations of ABT-199 (0-32 µM) in the absence and presence of either 0.125, 0.25 or 0.5 µM of S63845 for 72 hours. In the presence of 0.125 µM S63845, there was complete 100% cell killing resulting in 0% cell viability (S. Fig 5c – open circle) and the cells were sensitized to ABT-199 to >10-fold (S. Table 1) indicating maximum cell death can be achieved with lower drug concentrations, hence the fold-change in sensitivity could be higher. Similar data were obtained when the concentration of S63845 was increased to 0.25 µM (S. Fig 5c – open triangle & S. Table 1) and 0.5 µM S63845 (S Fig 5c – open diamond & S. Table 1).

Drug interaction analyses were conducted using the CompuSyn 1.0 software (ComboSyn Inc. NJ, USA) to generate Fa-CI isobologram plots. Points with CI < 1 indicates synergistic drug interaction. Drug interaction analyses indicated that the drug combinations demonstrated very strong and strong synergism at multiple doses of combinations of ABT-199 and S63845 (S. Fig 5d-f).

### Substantial inhibition of NPC cell proliferation driven by co-inhibition of MCL-1 and BCL-XL

Next, the NPC cell lines were treated with increasing doses of A-1331852 (0-32µM) and fixed doses of S63845 (0.5 µM, 1 µM or 2 µM) for 72 hours. In the HK-1 cells, the presence of 0.5 µM S63845, complete 100% cell killing was achieved resulting in 0% cell viability (Fig. 2a – open square) and S63845 sensitized the cells to A-1331852 by >15-fold (Table 2). Similar data were obtained when the concentration of S63845 was increased to 1 µM (Fig. 2a – open diamond & Table 2) and 2 µM (Fig. 2a – open triangle & Table 2).

**Fig. 2:**
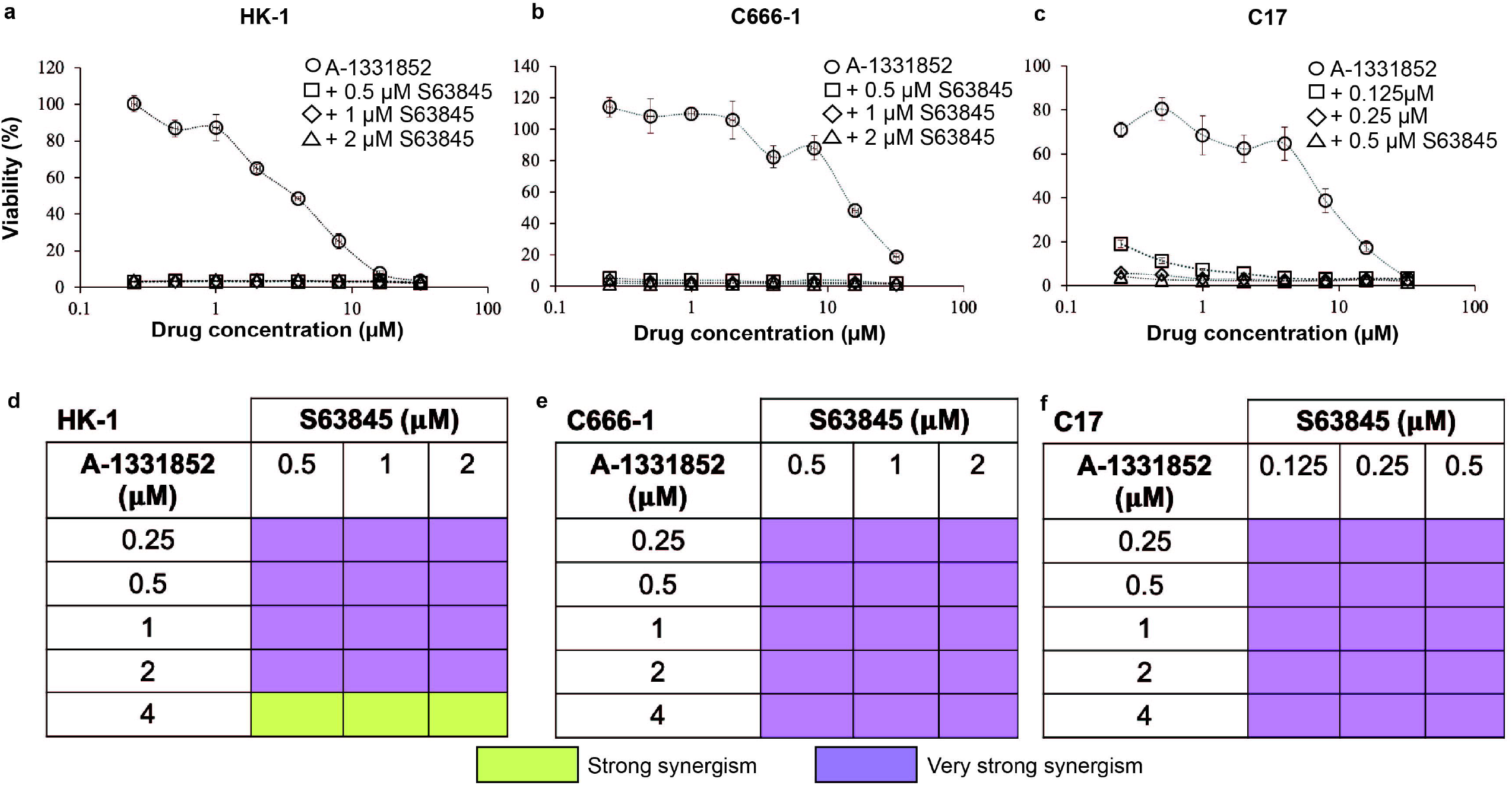
Co-inhibition of BCL-XL and MCL-1 using A-1331852 and S63845. NPC cell lines **(a)** HK-1; **(b)** C666-1 and **(c)** C17 cells were treated with increasing concentrations of A-1331852 (0–32 µM) in the presence and absence of S63845. Points represent mean SEM of four experiments. **(d-f)** Drug interaction analyses indicated that the combination of A-1331852 and S63845 demonstrated very strong and strong synergism at multiple doses of the drugs.

**Table 2:** Sensitization of NPC cell lines to A-1331852 by S63845 (fold sensitization). The IC_50_ values are concentrations of drug 1 **(bold)** that kill 50% of the cells surviving the shown concentrations of *drug 2 (italics)*. Fold sensitization: IC_50_ drug 1/ IC_50_ drug 2.

| **Cell line** | *S63845 (μM)* | **A1331852**<br>IC<sub>50</sub> ± SEM (μM) | **Fold sensitization by S63845** |
| --- | --- | --- | --- |
| **HK-1** | 0 | 3.8 ± 0.1 | - |
|  | 0.5 | <0.25* | 15.2 |
|  | 1 | <0.25* | 15.2 |
|  | 2 | <0.25* | 15.2 |
| **C666-1** | 0 | 15.5 ± 1.38 | - |
|  | 0.5 | <0.25* | 29.2 |
|  | 1 | <0.25* | 29.2 |
|  | 2 | <0.25* | 29.2 |
| **C17** | 0 | 5.9 ± 0.96 | - |
|  | 0.125 | <0.25* | 23.6 |
|  | 0.25 | <0.25* | 23.6 |
|  | 0.5 | <0.25* | 23.6 |

NOTE: \*Where the parent IC<sub>50</sub> was not calculable, the lower bound was used.

Similarly, in the C666-1 cells, combination with S63845 sensitized the cells to A-1331852 to all three concentrations tested. In the presence of 0.5 µM S63845, there was complete loss of the dose-dependent curve resulting in 0% cell viability (Fig. 2b-open circle) and S63845 sensitized the cells to A-1331852 by >29-fold (Table 2). Similar data were obtained when the concentration of S63845 was increased to 1 µM (Fig. 2b – open diamond & Table 2) and 2 µM (Fig. 2b – open triangle & Table 2).

As the C17 cells were sensitive to single agent S63845, we tested the sensitivity of the C17 cell lines to fixed concentrations of S63845 (below 1 µM) with increasing concentrations of A-1331852. At 0.125 µM, there was complete 100% cell killing resulting in 0% cell viability (Fig. 2c-open square). S63845 sensitized C17 cells to A-1331852, >23-fold (Table 2). Similar data were obtained when the concentration of S63845 was increased to 0.25 µM (Fig. 2c-open diamond & Table 2) and 0.5 µM (Fig. 2c -open triangle & Table 2).

Drug interaction analyses indicated that the drug combinations demonstrated very strong and strong synergism at multiple doses of A-1331852 and S63845 (Fig. 2d-f).

### S63845 sensitized spheroids to A-1331852 and *vice versa*

Next, combination of S63845 and A1331852 was tested on spheroids generated from the HK-1 cells. The HK-1 spheroids were treated with S63845 and A1331852 alone and in combination, first, over 3 days and later, over 10 days, to observe for emergence of resistance cells. The 3 days experiment findings demonstrated that the spheroids were insensitive to single agent treatment of S63845 (Fig. 3) and A-1331852 (Fig. 3) except at 2 µM concentration of each drug.

**Fig. 3:**
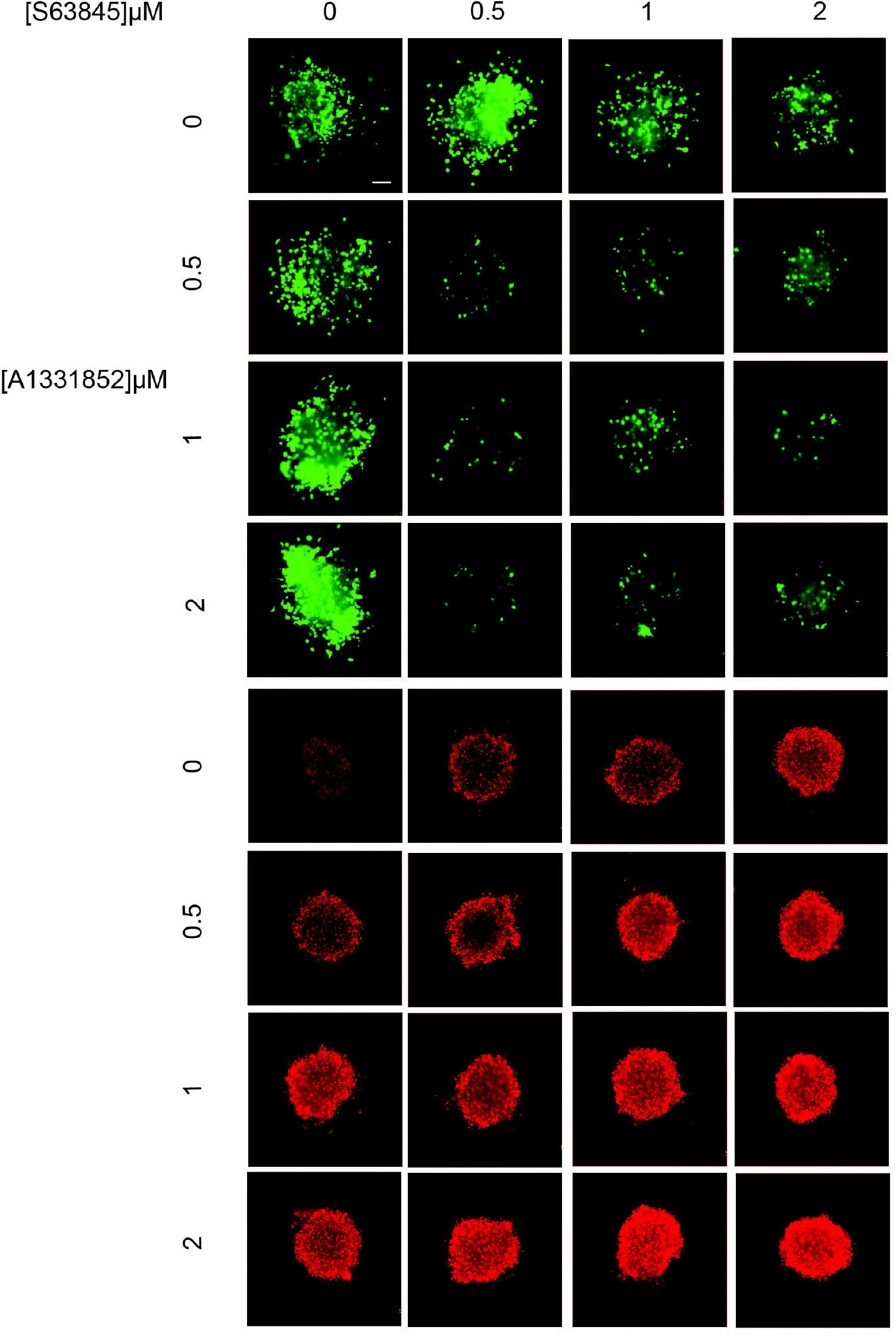
The effect of combination of S63845 and A-1331852 on the growth and invasion of HK-1 spheroids over three days. The spheroids were treated with single agents S63845 and A-1331852 and combination of both over three days at the indicated concentrations, n=2–3 spheroids per combination. Cell viability was determined using the live/dead assay (Viable cells: stained green by Calcein-AM; Dead cells: stained red by Ethidium-homodimer I). Size bar: 200 µm.

There was obvious sensitization of the spheroids to A-1331852 by S63845 at concentration of S63845 as low as 0.5 µM. Similarly, A-1331852 was also able to sensitize the spheroids to S63845 at 0.5 µM, which manifested as reduced spheroid invasion (Fig. 3). Further increase of concentrations of both inhibitors, reduced cell invasion in a dose-dependent manner (Fig. 3). This was evident with the increase of red fluorescence intensity indicating more dead cells as the combination concentrations increased. Over 10 days, treatment with combination of S63845 and A-1331852 resulted in greater reduction of spheroid growth and invasion. Moreover, there were no emergence of green fluorescence or in other words emergence of viable cells indicating there were no rapid generation of resistance cells over 10 days (S. Fig 6).

Similar data were obtained when the spheroids generated from the C17 cells were tested to combination of S63845 and A-1331845. There was obvious sensitization of the spheroids to A-1331852 by S63845 at concentration of S63845 as low as 0.5 µM. Similarly, A-1331852 was also able to sensitize the spheroids to S63845 at 0.5 µM, which manifested as reduced spheroid invasion (S. Fig 7). Further increase of concentrations of both inhibitors, reduced cell invasion in a dose-dependent manner. This was evident with the increase of red fluorescence intensity indicating more dead cells as the combination concentrations increased (S. Fig 7). There was no emergence of resistance cells over 10 days (S. Fig 8).

Collectively, our data demonstrate that there was a greater response of the NPC cell lines to co-inhibition of MCL-1 and BCL-XL compared to co-inhibition of MCL-1 and BCL-2, suggesting that co-inhibition of MCL-1 and BCL-XL are better therapeutic targets for killing NPC cells.

### *BFL-1* deleted NPC cells sensitized to BCL-XL selective inhibitor A-1331852

The C666-1 cells expressed undetectable levels of *BFL-1* (S. Fig 3b). Hence, to access the role of *BFL-1* in cell survival, the gene was deleted in the HK1 cells. The HK-1 cells were transfected with two independent single-guide RNAs (sgRNAs) targeting different regions of the human *BFL-1* gene (hereafter the sgRNAs will be referred to as sg*BFL-1*#1 and sg*BFL-1*#2). The experimental design is shown in Fig. 4a. *BFL-1* mutagenesis was verified by conventional Sanger DNA sequencing and gene expression of *BFL-1* in the parental and *BFL-1* deleted cells were verified by qPCR. The sg*BFL-1*#1 generated higher number of InDels (insertions and deletions of bases in the DNA) compared to sg*BFL-1*#2 (Fig. 4b). The sg*BFL-1*#1 and sg*BFL-1*#2 resulted in ∼98% and 80% reduction in *BFL-1* expression, respectively (Fig. 4c).

**Fig. 4:**
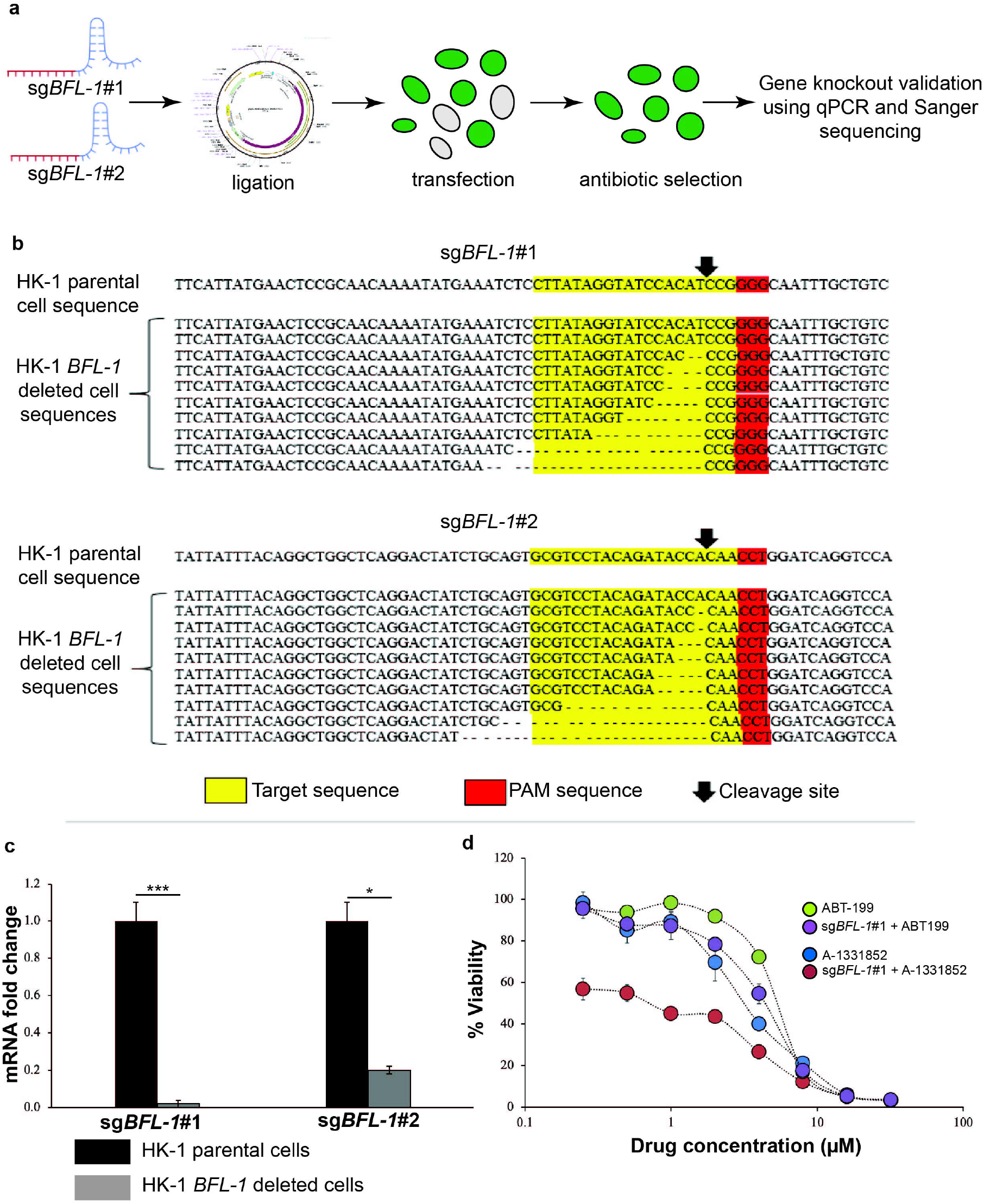
HK-1 *BFL-1* deleted cells were sensitive to treatment of BCL-XL selective inhibitor A-1331852. **(a)** Experimental design to generate the HK-1 *BFL-1* deleted cell lines using the CRISPR/Cas9 technique. **(b)** Read sequences of the HK-1 sg*BFL-1*#1 and HK-1 sg*BFL-1*#2 cells. Parental *BFL-1* sequence is shown on the top. Most of the clones tested showed deletions at the expected cleavage sites (arrow). The dashed lines indicate the InDels. The target sequence is highlighted in yellow while the PAM sequence is highlighted in red. **(c)** qPCR validation *BFL-1* gene deletion in the HK-1 cells. *BFL-1* expression levels were normalized to parental cells. Bars indicate mean SEM of three independent experiments. Statistically significant differences in *BFL-1* expression between the parental cell line and the *BFL-1* knockout cells are shown as ***p ≤ 0.001 or *p < 0.05 determined by two-tailed paired T-test.**(d)** The sensitivity of the HK-1 parental cell line and the HK-1 sg*BFL-1*#1 cells were tested to tested to single agent activity of either ABT-199 or A-1331852 (0-32 µM). Points represent mean SEM of four experiments.

Given that the sg*BFL-1*#1 demonstrated more profound reduction of *BFL-1* expression, the sg*BFL-1*#1 cells were treated with increasing concentrations of either ABT-199 or A1331852 to access whether *BFL-1* deletion sensitized NPC cells to these inhibitors. The HK-1 parental cells were resistant to single agent treatment of ABT-199 and A-1331852 (Fig. 4d). The sg*BFL-1*#1 cells were weakly sensitized to ABT-199 (Fig. 4d – purple circle & S. Table 2). However, the sg*BFL-1*#1 cells were sensitized to A-1331852 by ∼5-fold (Fig. 4d – red circle & S. Table 2), indicating that the BFL-1 and BCL-XL may be important for NPC cell survival.

## Discussion

Given that different cell population are addicted to different anti-apoptotic protein(s) for survival, it is crucial to determine the anti-apoptotic proteins that NPC cells depend for survival. We combined the CRISPR/Cas9 technique and BH3 mimetics to delineate the individual contributions of the anti-apoptotic proteins for NPC survival. BH3 mimetics, given their highly selective inhibition of the anti-apoptotic proteins, provide a chemical toolkit to parse the individual contributions of the anti-apoptotic proteins for cancer cell survival.

In our hands, deletion of MCL-1 alone did not kill the HK-1 cells, implying that other anti-apoptotic proteins may be compensating for the loss of MCL-1. Similarly, all three NPC cell lines tested were insensitive to single agent treatment of ABT-199 and A-1331852. Except for C17, none of the other cell lines responded to single agent treatment of S63845, indicating that these cell lines do not depend solely on MCL-1 for survival. Although the C17 cells responded to single agent treatment of S63845, combination treatment with either ABT-199 or A-1331852 with concentrations of S63845 of < 1 µM resulted in synergy, demonstrating that co-inhibition with either BCL-2 or BCL-XL is still needed to achieve maximal cell killing at lower concentrations. Collectively, these findings reveal that NPC cells depend on more than one anti-apoptotic protein for survival and targeting them appropriately can result in significant cell death.

The HK-1 sgMCL-1#2 cells were weakly sensitized to ABT-199 but were sensitized to A-1331852 indicating that MCL-1 and BCL-XL may be important for NPC cell survival. However, gradual increase of S63845 concentrations, sensitized all NPC cell lines tested to ABT-199. The effect of co-inhibition MCL-1 and BCL-XL with S63845 and A-1331852 were more profound as all three NPC cell lines were instantly killed at nanomolar drug combination concentrations. It appears that combination of BH3 mimetics were able to induce more severe phenotype than was observed with the CRISPR/Cas9-manipulated cells. Although complete inhibition of *MCL-1* was achieved with sgMCL-1#2, the *MCL-1* manipulated cells used were bulk cells. The manipulated cells did not survive single cell clonal expansion, despite multiple attempts. Hence, the mild sensitivity of the *MCL-1* manipulated cells to the BH3 mimetics could be due to the heterogenous cell population.

The sensitization of the HK-1 and C666-1 to ABT-199 increased to 16-and 12-fold, respectively at concentration of 1 µM of S63845 and the sensitization of the cell lines to ABT-199 improved at 2 µM of S63845. Both cell lines were only modestly sensitized to ABT-199 at concentrations < 1 µM of S63845. The C17 cells were sensitized to ABT-199 at nanomolar concentrations of S63845. Compared to co-inhibition of MCL-1 and BCL-2, all three cell lines responded profoundly to co-inhibition of BCL-XL and MCL-1. The HK-1 and C666-1 cells were sensitized to A-1331845 by 15-and 29-fold, respectively at concentration of S63845 at 0.5 µM. Cells attained 0% viability at nanomolar combination doses. Similarly, in the C17 cells, the cells were sensitized to A-1331845 at concentration of S63845 as low as 0.125 µM, indicating that BCL-XL and MCL-1 are crucial for NPC cell survival. The sensitization obtained in the monolayer culture were analogous with findings obtained from the spheroid studies. Spheroids generated from the HK-1 and C17 cells were sensitized to A-1331852 by S63845 and *vice versa*. Moreover, combination of S63845 and A-1331852 inhibited spheroid growth and invasion at low doses, indicating that the combination may be effective *in vivo*. In accordance with our study findings, many solid tumours are dependent on BCL-XL and MCL-1 for survival [11]. However, targeting BCL-XL and MCL-1 in the clinic may be cumbersome. Inhibition of BCL-XL may result in thrombocytopenia [12, 13] and co-inhibition of BCL-XL and MCL-1 can result in fatal hepatotoxicity [14]. There are a few strategies to overcome this issue. Our data shows that cell killing can be achieved at very low doses of S63845 and A-1331852, which may not be sufficient to cause thrombocytopenia. Another strategy would be to co-inhibit BCL-2 and MCL-1. Although the effect of inhibiting BCL-2 was not as pronounced as inhibiting BCL-XL, we still observed synergy with combination of ABT-199 and S63845 in all NPC cell lines tested. Hence targeting BCL-2 and MCL-1 could be an alternative treatment strategy. In a separate study, co-inhibition of BCL-2 and MCL-1 demonstrated synergistic effects in inducing killing of NPC cells *in vitro* and *in vivo* [15]. Another strategy, although still preliminary, is to co-target BCL-XL and BFL-1.

*BFL-1* is a direct target gene of the NFκB signaling pathway [16] which largely drives NPC pathogenesis [17]. However, the role of BFL-1 for NPC survival was not reported previously. In our hands, the sg*BFL-1*#1 produced more InDels and reduced the exogenous *BFL-1* expression more profoundly compared to sg*BFL-1*#2. Despite achieving complete reduction of *BFL-1* expression with sg*BFL-1*#1, no notable reduction of cell viability was observed in the sg*BFL-1*#1 deleted cells suggesting that inhibition of *BFL-1* alone is not sufficient to activate cell death in the NPC cells. The sg*BFL-1*#1 deleted cells were modestly sensitized to ABT-199 but were sensitized to A-1331852 treatment. This is the first study which demonstrates the involvement of BFL-1 for NPC survival. Given that both BFL-1 and BCL-XL are direct targets of NFκB [18], these anti-apoptotic proteins may have important implications for NPC cell survival. Therefore, to omit the possibility that this is a cell-type effect and to further interrogate their functional roles for NPC cell survival, similar experiments need to be conducted in other NPC cell lines and models. Moreover, testing the sensitivity of the cells to BFL-1 selective inhibitor would be useful but thus far there are no selective inhibitors reported for BFL-1.

Future work should focus on using mouse models to access the toxicity profiles of the drug combinations. Moreover, expression of the anti-apoptotic proteins should be determined in patient-derived tissues. Nonetheless, with the findings put forward, MCL-1 and BCL-XL are crucial for NPC cell survival and targeting these proteins at appropriate doses will yield maximal cell killing. Moreover, the possibility of involvement of BFL-1 for cell survival may open new treatment avenues for NPC.

## Funding

This study was funded by the Fundamental Research Grant Scheme (FRGS), Ministry of Education, Malaysia (Grant number: 203/PBIOLOGI/601228), Universiti Sains Malaysia Research University (RU) grant (Grant number: 1001/PBIOLOGI/8012268) and Loreal Malaysia. We would like to thank Professor Dr. George Sai Wah Tsao (University of Hong Kong, Pokfulam, Hong Kong, China) for providing the NPC cell lines HK-1 and C17.

## Conflicts of interest

The authors declare that there are no conflicts of interest.

## Availability of data and material

All data are available from the corresponding author upon reasonable request.

## Authors’ contributions

SFAR: performed the experiments, data analysis, writing – original draft; AA: helped SFAR with experiments which involved the CRISPR/Cas9 technique; K-WL: resources; GA: resources; NM-K: conceptualization, data analysis, resources, writing – original draft, funding acquisition.

## Notes

### Competing Interest Statement

The authors have declared no competing interest.

