## Supplementary data for "Dual inhibition of anti-apoptotic proteins BCL-XL and MCL-1 enhances cytotoxicity of Nasopharyngeal carcinoma cells"

**Supplementary Information**

**Materials and Methods**

**Cell culture**

The human NPC cell lines, HK-1 and C666-1 were obtained from the Molecular Pathology Unit, Institute for Medical Research, Selangor, Malaysia after obtaining permission from Professor George Tsao (University of Hong Kong) for the HK-1 cells and Professor KW Lo (Chinese University of Hong Kong) for the C666-1 cells. The C17 cells were obtained from Universiti Malaya, Malaysia after obtaining permission from Professor George Tsao (University of Hong Kong). The HK-1 and C666-1 cell lines were cultured in RPMI 1640 medium supplemented with 10% heated fetal bovine serum (FBS). Additionally, 10% (v/v) Glutamax was added to the RPMI complete medium that was used to grow the C666-1 cells. NPC cell lines HK-1 and C666-1 were authenticated using the AmpFISTR profiling. The C17 cells were cultured in RPMI medium supplemented with 10% heated fetal bovine serum and Rho inhibitor (4 µM in 500 ml medium) (Enzo Life Sciences, NY, USA). The C17 cells were authenticated using the short tandem repeat (STR) analysis (Yip *et al.,* 2018). Cells were sub-cultured when required.

**RNA extraction**

Cells were trypsinized and centrifuged at 500 g (Sigma 3-16PK, rotor number:12171) for 5 minutes. Cell pellets were washed with 1X PBS and re-centrifuged at the same speed for another 5 minutes. The PBS (supernatant) was removed and 1 ml of TRIzol (Thermo Fisher Scientific, MA, USA) was added to the cell pellet and re-suspended several times for the cells to undergo complete lysis. The mixture was incubated at room temperature for 5 minutes. Next, 0.2 ml of chloroform was added to the mixture and the tubes were inverted vigorously for about 15 seconds. The tubes were then incubated at room temperature for 5 minutes. After 5 minutes, the samples were centrifuged at 12000 g (Sigma 3-16PK, rotor number:12171) for 15 minutes at 4^o^C. Phase separation was observed where the uppermost layer was composed of RNA, the middle layer was occupied by DNA and the bottom layer occupied by proteins. Approximately, 0.45 ml of the top-most layer (RNA) was transferred into a new 1.5 ml tube. Equal volume of isopropanol was added to each tube and mixed well. The mixture was then incubated at room temperature for 5 minutes prior to centrifugation at 12000 g for 5 minutes. The supernatant was removed and the pellet was re-suspended with 1 ml of ice cold 75% ethanol and re-pelleted at 7500 g for 5 minutes at 4^0^C. Next, the RNA pellets were air dried at room temperature for 3-5 minutes and dissolved in 30 µl nuclease-free water (Qiagen, Hilden, Germany). The RNA yield and purity of the total RNA extracted, were assessed using Nanodrop 1000^TM^ (Thermo Fisher Scientific, MA, USA). Absorbance of the samples were measured at 260 nm and 280 nm wavelengths and the 260/280 ratio was used to assess RNA purity. The RNA samples were considered pure when the 260/280 ratio fall between 1.8-2.0. Lower ratio could indicate that the RNA samples are contaminated with either proteins, phenol or other contaminants. The RNA samples were stored at -80^0^C until further use.

**Custom RT^2^ Profiler PCR Array**

*Prior to knocking-out the anti-apoptotic genes MCL-1 and BFL-1 using the CRISPR/Cas9 technique, the mRNA levels of all anti-apoptotic genes, BCL-2, BCL-XL, BCL-w, MCL-1 and BFL-1 in the NPC cell lines were determined using a Custom RT^2^ Human Apoptosis Profiler PCR array.*

The custom human apoptosis profiler array which was used in this study was designed by Qiagen, Hilden, Germany. The profiler was in a 96 well plate format and contained probes to detect and amplify 20 customized pro- and anti-apoptotic genes (see Appendix 1 for the list of customized pro- and anti-apoptotic genes for this study). The RT^2^ First Strand Kit consists of a genomic DNA elimination mix and a reverse-transcription mix. The genomic DNA elimination mix was prepared for each RNA sample. The RNA samples extracted earlier (see 3.2.3) were mixed gently with the DNA elimination mix. This is to remove any residual DNA contaminations in the RNA samples. The samples with the elimination mix were incubated at 42^0^C for 5 minutes then placed in ice immediately for 1 minute. Next, the reverse- transcription mix was prepared. Ten microliters of the reverse-transcription mix were added to each tube containing the DNA elimination mix and incubated at 42^0^C for 15 minutes. The reaction was stopped immediately by incubating the tubes at 95^0^C for 5 minutes. Ninety-one microliters of RNase-free water (Qiagen, Hilden, Germany) was added to each reaction and mixed well by pipetting up and down several times. The cDNA samples were stored at -20^o^C for long term storage. The PCR components mix was prepared and the cDNA samples were added to the mix. Twenty-five microliters of the mix were added to each well of the customized RT^2^ PCR array 96 well plate. The plate was centrifuged for 1 min at 1000 g (Kubota 2800, rotor number: RMP-23) to remove bubbles. The real-time cycler conditions were as follows (95^0^C 10 min X 1, [95^0^C 15s, 60^0^C 1 min] x 40) (Applied Biosystem, Thermo Fisher Scientific, New York, USA).

**Two-dimensional (2D) Cytotoxicity Assay conducted with parental NPC cell lines HK-1, C666-1 and C17**

The HK-1 cells were seeded at a density 3500 cells/well, the C666-1 cells were seeded at a density of 4000 cells/well and the C17 cells were seeded at a density of 6000 cells/well in 96 well plates (Corning, New York, USA). The cells were left to attach for 5-6 hours in a humidified incubator at 37^0^C with 5% C0_2_. Cells were viewed under the inverted phase contrast microscope (Olympus CKX41, Tokyo Japan) to ensure cell attachment prior to adding the BCL-2 selective family inhibitors. The first three columns of the 96-well plate, only contained cells (positive controls) while the last column only contained only media (negative control). The columns in between were treated with different concentrations of BH3 mimetics. First, the cells were treated with single agents of the BH3 mimetics - ABT-199 which inhibits anti-apoptotic protein BCL-2, S63845 which selectively inhibits anti-apoptotic protein MCL-1 and A-1331852 which selectively inhibits anti-apoptotic protein BCL-XL, in two-fold increasing doses from 0.25 µM to 32 µM. All BH3 mimetics were purchased from MedChem Express, USA.

Subsequently, the cells were treated with combinations of two inhibitors where a fixed dose of one selective inhibitor was added to increasing concentrations of another selective inhibitor. As for the HK-1 and C666-1 NPC cells, a fixed dose of S63845 (0.5 µM, 1 µM or 2 µM) was added to two-fold increasing doses (0.25-32 µM) of either ABT-199 or A-1331852. As for the C17 NPC cells, a fixed dose of S63845 (0.125 µM, 0.25 µM or 0.5 µM) was added to two-fold increasing doses (0.25-32 µM) of either ABT-199 or A1331852.

**Drug Combination Analyses**

Synergistic interaction between the drug combination were analyzed using the CalcuSyn 2.11 software (Biosoft, Cambridge, UK). The software was employed to produce the combination index (CI) values, which is a statistical measure of synergy. CI values < 1 indicate synergy, = 1 indicate an additive effect and > 1 indicate antagonism. C1 values nearing 0 indicate strong synergy.

**Knocking-out anti-apoptotic genes *MCL-1* and *BFL-1* in NPC cell lines HK-1 and C666-1 using the CRISPR/Cas9 technique**

*Anti-apoptotic genes MCL-1 and BFL-1 were knocked out to determine the contribution of this gene to NPC cell survival.*

***Designing of the single-guide RNAs (sgRNAs)***

The CHOP CHOP design software was employed to design sgRNAs (<https://chopchop.cbu.uib.no/>) for the targeted genes, *MCL-1* and *BFL-1.* Two sgRNAs were designed. The sgRNAs were cloned into an expression vector plasmid via BbSI (Vivantis Technologies, Setia Alam, Malaysia) digestion. The sgRNAs oligo design included a 4-bp overhang for the forward (CACC – see highlighted region in the sequences below) and complementary reverse (CAAA) to enable cloning using BbSI digestion. The customized oligos were synthesized by Intergrated DNA Technologies (IDT), Singapore.

***MCL-1* gRNA oligo sequences (5^’^- 3^’^)**:

1^st^: Forward oligo: 5^’^ – CACC GGGAGGGCGACTTTTGGCTA – 3^’^

Reverse oligo: 5^’^ – AAAC TAGCCAAAAGTCGCCCTCCCC – 3^’^

2^nd^: Forward oligo: 5^’^ – CACC GGAGCTGGACGGGTACGAGC – 3^’^

Reverse oligo: 5^’^ – AAAC GCTCGTACCCGTCCAGCTCCC – 3^’^

***BFL-1* gRNA oligo sequences (5^’^- 3^’^)**:

1^st^: Forward oligo: 5^’^ – CACC CTTATAGGTATCCACATCCG – 3^’^

Reverse oligo: 5^’^ – AAAC CGGATGTGGATACCTATAAGC – 3^’^

2^nd^: Forward oligo: 5^’^ – CACC GTCCTACAGATACCACAACC – 3^’^

Reverse oligo: 5^’^ – AAAC GGTTGTGGTATCTGTAGGACC – 3^’^

***Cloning of MCL-1 and BFL-1 sgRNAs into plasmid vector***

The sgRNA oligos were termed as sg*MCL-1*#1, sg*MCL-1*#2, sg*BFL-1*#1 and sg*BFL-1*#2. These oligos were individually cloned into an ‘all-in-one’ one vector system by restriction digestion and ligation. Cloning of sgRNA into plasmid vector comprises of five steps which are (1) linearization of the plasmid vector backbone (Px549 plasmid) (S. Fig 1), (2) phosphorylation and annealing of the sgRNA oligos, (3) ligation of the linearized plasmid with annealed sgRNA oligos, (4) transformation of the ligated plasmid into competent cells (Rubidium Chloride Competent Cell), and (5) sequence verification of the sgRNAs. Details of the steps are as below:

***Step 1: Linearization of the PX459 plasmid vector backbone***

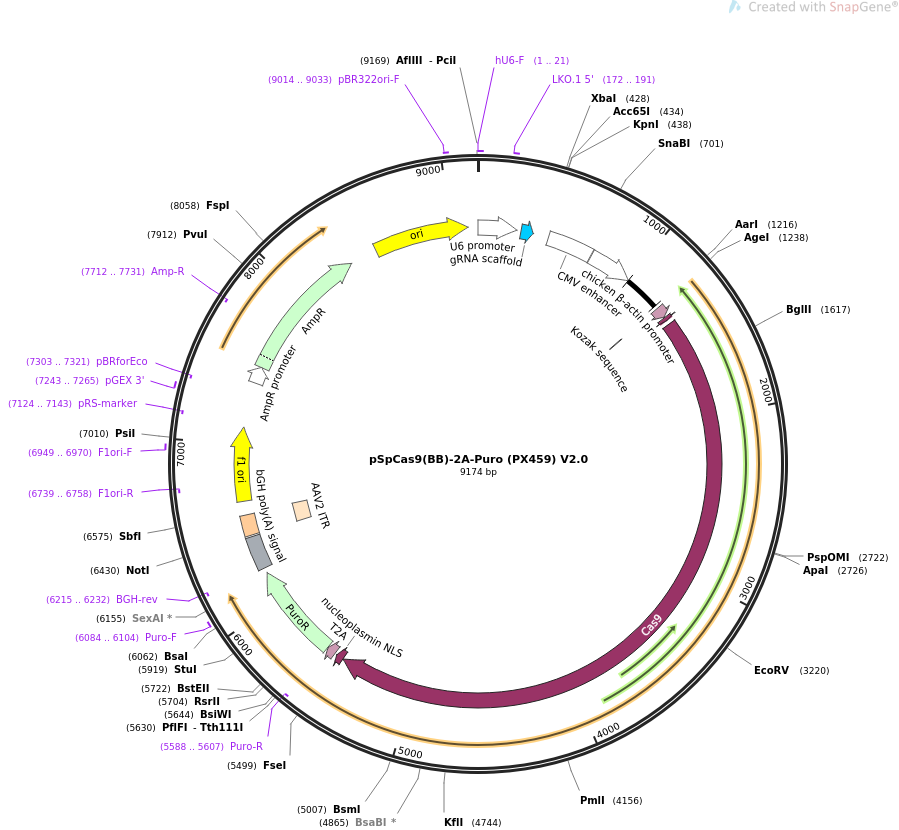

Figure source: Addgene (https://www.addgene.com)

**S. Fig 1:** Plasmid map of PX459 vector. Red box indicates the restriction enzyme cut site. The sgRNA of the target sequence was inserted in the BbSI cut site.

a. Plasmid linearization was performed to allow ligation with either sg*MCL-1*#1, sg*MCL-1*#2, sg*BFL-1*#1or sg*BFL-1*#2, sequences. In order to linearize the Px459 plasmid (62988, Addgene, MA, USA), the following reagents were mixed well and incubated at 37^0^C overnight:

4 µl plasmid

1 µl Bpil (Fast Digest BbSI)

3 µl Buffer G

Top up double distilled water to final volume of 20 µl

b. After an overnight digestion, the digestion mixture was run on 1% agarose gel. A thin band was observed for the digested plasmid (9175kb). The digested plasmid was purified using the Wizard® SV Gel and the PCR Clean-up System (Promega, Wisconsin, USA) according to the manufacturer’s protocol. The concentration and purity of the purified plasmids were evaluated using Nanodrop 1000^TM^ (Thermo Fisher, New York, USA). The purified plasmid stocks were stored at -20 ^0^C.

***Step 2: Phosphorylation and annealing of the MCL-1 or BFL-1 sgRNA oligos***

c. The following reaction mixture was prepared to anneal and phosphorylate either the sg*MCL-1*#1, sg*MCL-1*#2, sg*BFL-1*#1, or sg*BFL-1*#2 oligos:

1 µl forward oligo (100 µM)

1 µl reverse oligo (100 µM)

1 µl 10X T4 ligase buffer

1 µl 10X T4 ligase buffer

6 µl double distilled water

The reaction was placed in a thermal cycler (BioRad, California, USA) and run using the following parameters: 37^0^C for 30 minutes; 95^0^C for 5 minutes; ramp down to 25^0^C at 5^0^C min^-1^. The annealed oligos were then diluted at 1:200 ratios with double distilled water.

***Step 3: Ligation of the linearized plasmid with the annealed MCL-1 or BFL-1 sgRNA oligos***

d. In order to ligate the sgRNA into the linearized plasmid backbone, the following reaction was set up. A

reaction excluding the sgRNA was included to confirm the efficiency of the digestion and ligation:

**X** µl (50 ng) of plasmid

The mixture was incubated at room temperature for 30 minutes.

1 µl diluted annealed oligo

1 µl T4 DNA Ligase

2 µl 10X T4 ligase buffer

Top up double distilled water to final volume of 10 µl

***Step 4: Transformation of the ligated plasmid into competent cells (Rubidium Chloride Competent Cell)***

e. The ligated oligo-plasmid was then transformed into the competent cells (Rubidium Chloride Competent Cell) using the heat shock method to allow selection of successfully ligated oligos with the plasmid backbone. Five microliters of the ligation reaction were added to 25 µl of competent cells and incubated in ice for 30 minutes. After 30 minutes, the cells were heat shocked at 42^0^C for 45 seconds. The cells were immediately incubated in ice for 3 minutes to stop the heat shock reaction.
f. The transformed cells were activated by adding 250 µl of sterile LB broth (First Base, Singapore) and mixed at 37^0^C for 45 minutes at 200 rpm, allowing the bacterium to recover and express ampicillin resistance marker encoded by the plasmid.
g. One hundred microliters of the culture were plated onto LB agar plates supplemented with 100 µg/ml of ampicillin (Nacalai Tesque, Kyoto, Japan). Plates were incubated at 37^0^C, overnight. Empty competent cell (without plasmid) was used as negative control to verify successful transformation.
h. Three to four colonies were picked per gRNA and inoculated into sterile LB broth supplemented with 100 µg/ml of ampicillin and cultured overnight at 37^0^C at 200rpm. Glycerol stocks were made for each colony by mixing the overnight LB culture with 50% glycerol at 1:1 ratio and stored at -80^0^C.

***Step 5: Sequence verification of the MCL-1 or BFL-1 sgRNAs by Sanger sequencing***

The gRNA sequences were verified with U6 primers (forward primer: GAGGGCTATTTCCATGATTCC, reverse primer: GCAACACACAACATCTCCA). Vectors were purified from the glycerol stocks in LB broth using the GF-1 Plasmid Extraction Kit (Vivantis, Subang Jaya, Selangor). Sequence of selected colonies from each gRNA was sequenced with U6 forward primer via Sanger sequencing by First Base, Singapore.

***Transfection of the MCL-1 or BFL-1 sgRNAs into Nasopharyngeal Carcinoma (NPC) Cell Lines***

The *MCL-1* sgRNAs and the *BFL-1* sgRNAs were introduced into NPC cell line HK-1 cells via transfection. Lipofectamine 2000 (Thermo Fisher Scientific, MA, USA) was used as transfection reagent due to its high transfection rates. Since the vector Px459 housed Puromycin as the selection marker, the optimum concentration of Puromycin for selection was first determined. The following detail the steps involved in the transfection of either the *MCL-1* or *BFL-1* sgRNAs into the NPC cell lines.

***Step 1: Determination of optimum concentration of Puromycin for selection of manipulated cells***

a. The HK-1 cells were seeded into individual 96 well plates at 2 × 10^4^ cells per well. The plates were incubated in a humidified incubator at 37^0^C with 5% CO_2_ overnight.
b. The following day, medium was aspirated from each well and the cells were washed with 200 µl of 1X PBS. An independent dilution of Puromycin in 200 µl of complete media was added to both the cell lines. Puromycin was titrated within the range of 1-7 µg/ml. The wells without Puromycin served as controls. The plates were incubated in a humidified incubator at 37^o^C with 5% CO_2_ and monitored at 24-hour intervals for 5 days.
c. After 5 days, surviving cells were counted using trypan blue (see section 3.2.2) to determine the lowest concentration of Puromycin required to kill all non-transfected cells. The lowest concentration of Puromycin obtained, was used as the optimal concentration for selection of transfected cells.

***Step 2: Transfection***

d. The NPC cells HK-1 and C666-1 were seeded in individual twenty-four well plates at 1 × 10^5^ cells per well. The cells were plated in an antibiotic-free growth medium. The plates were incubated overnight in a humidified incubator at 37^0^C with 5% CO_2_ until the cells reached 80% cell confluency.
e. On the day of transfection, 3 µl Lipofectamine 2000 (Thermo Fisher Scientific, MA USA) was added to 300 µl of serum reduced media (called ‘tube A’). Three micrograms of DNA of vector carrying either the *MCL-1* or *BFL-1* sgRNAs were added to 300 µl of serum reduced media (called ‘tube B’). Both tubes were incubated at room temperature for 30 minutes.
f. After 30 minutes, content of tube A was transferred to tube B and mixed well. Tube B was incubated at room temperature for another 30 minutes to form the transfection complex (transfection reagent + DNA).
g. Media were aspirated from each well of the twenty-four well plates and serum reduced media was added to each well. The transfection complex mixtures were added to each well ‘dropwise’. The twenty-four well plates were incubated at 37^0^C with 5% CO_2_ for 72 hours.\
h. After 72 hours, the transfected cells were washed with 2 ml of 1X PBS. In order to select for transfected cells, 1 ml of serum reduced media supplemented with 1ug/ml Puromycin was added to each well. The plates were incubated in a humidified incubator at 37^0^C with 5% CO_2_. After 48 hours, the selection was stopped and complete media was added into each well. Once each well reach about 80% confluency, the cells were expanded in order to extract DNA and RNA to validate the gene knockouts via qPCR and sequencing.

***Validation of the MCL-1 and BFL-1 knock-outs***

*Following the transfection of the vector carrying either the MCL-1 or BFL-1 sgRNAs, into the NPC cells, the following validation steps were taken to ensure that the genes of interest were indeed knocked-out in the NPC cell line HK-1.*

***Step 1: DNA extraction from potential MCL-1 or BFL-1 knock-out HK-1 cells***

a. Manipulated HK-1 cells were trypsinized and transferred into a 15 ml tube and centrifuged at 1500 rpm for 10 minutes (Kubota 2010, rotor number: RS-240). The supernatant was removed and the pellet was washed with 1X PBS.
b. The cell suspension was pelleted again at 1500 rpm for 10 minutes. The supernatant was removed and 1 ml of the DNA digestion cocktail was added to the cell pellet. The cells were incubated at 37^0^C overnight.
c. The following day, phenol-chloroform was added at 1:1 ratio to the incubated cell solution to purify the DNA. The mixture was centrifuged at 14,000 rpm (Kubota 2010, rotor number: RS-240) for 10 minutes at 4^0^C.
d. The top layer (clear solution) containing the DNA was transferred into a new 1.5 ml tube and mixed with absolute ethanol (99%) at 1:1 ratio.
e. The mixture was centrifuged at 14,000 rpm (Kubota 2010, rotor number: RS-240) for 10 minutes at 4^0^C. The supernatant was removed and the pelleted DNA was washed with 70% ethanol and re-suspended with 30 µl of T_10_E_0.1_ buffer.
f. The concentration and purity of the extracted DNA were evaluated using Nanodrop 1000^TM^ (Thermo Fisher, New York, USA). The DNA stocks were stored at -20 ^0^C.

***Step 2: Confirmation of either MCL-1 or BFL-1 gene knock-outs using PCR***

a. Primers were designed for each gRNA of both the genes to screen for the genomic knockouts. The primers recognized sequences approximately ~100-200 bp within the CRISPR cut site. Wild type population for each cell line used as control for the PCR reaction.

***Primers to screen for MCL-1* gRNA knockout (5^’^- 3^’^)**:

1^st^: Forward oligo: 5^’^ – GTT TGG CCT CAA AAG AAA CG – 3^’^

Reverse oligo: 5^’^ – CTC TCT ATC CCC CTC CCC – 3^’^

2^nd^: Forward oligo: 5^’^ – CCG CTT GAG GAG ATG GAA G – 3^’^

Reverse oligo: 5^’^ – TAT TGT GGT CAT GCC TGC CCG – 3^’^

***Primers to screen for BFL-1* gRNA knockout (5^’^- 3^’^)**:

1^st^: Forward oligo: 5^’^ – AGC CTC CGT TTT GCC TTA TC– 3^’^

Reverse oligo: 5^’^ – GAA GGG GTC AAT TAC TAC GG – 3^’^

2^nd^: Forward oligo: 5^’^ – TCT CAG CAC ATT GCC TCA AC – 3^’^

Reverse oligo: 5^’^ – TCG TTT TGC AGG TCT CAC GA – 3^’^

b. During PCR assay, *Taq* polymerase was used to generate 3’A overhangs for TA cloning. The *Taq*

polymerase was used to generate 3’A overhangs for TA cloning. The PCR products were then purified using Wizard ® SV Gel and PCR Clean-up System (Promega, Wisconsin, USA).

c. The purified PCR products were then ligated into pGEM-T Easy Vector (A3600, Promega) (S. Fig 2). The ligation reaction mixture includes:

1 µl (50 ng) pGEM-T Easy Vector

4 µl Purified PCR product

1 µl T4 DNA Ligase

1 µl 10X T4 ligase buffer

3 µl Nuclease-free water

The ligation mixture was incubated at 4^0^C overnight. Next, 2 µl of the ligation mixture were transformed into competent cells according to the transformation protocol. The transformation culture was plated onto LB-ampicillin/IPTG/X-Gal plates. The plate was incubated overnight at 37^0^C. Successful cloning of an insert would interrupt the coding sequence of β-galactosidase encoded by the *lacZ* gene and the colonies would appear white. Unsuccessful cloning or self-ligation of the vector would produce blue colonies. The subsequent day, several white colonies were selected for colony PCR. Colony PCR was performed to determine the presence of insert DNA in the vector. Colonies that contained the correct insert sizes were verified by gel electrophoresis and later were grown in LB-ampicillin liquid media overnight at 37^0^C. The plasmids were isolated from the bacterial culture and sent for Sanger sequencing (First Base, Singapore) to verify the genomic knockout sequences. The mutated DNAs of the colonies were compared to the parental sequence to check for presence of indels and affirm the knockouts.

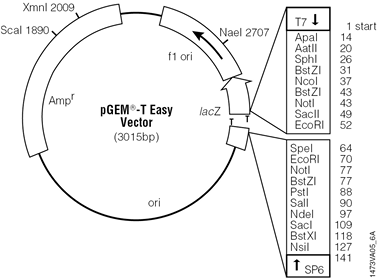

Figure source: Promega (https://worldwide.promega.com)

**S. Fig 2:** pGEM-T Easy Vector gene map. Red box indicates the site where PCR product were ligated. This location has the T overhang.

***Step 3: RNA Extraction from potential knock-out cells and qRT-PCR***

a. RNA was extracted and quantified from the *MCL-1* or *BFL-1* knock-out cells. Complementary DNA (cDNA) synthesis was performed using the iScript Reverse Transcription Supermix (Bio-Rad, California, USA). A total of one microgram of RNA was used in 20 μl reactions. The reaction set-up was as listed below:

4 μl iScript RT Supermix

**X** μl RNA template (1μg)

Top up RNase free water to final volume of 20 µl

The reaction mix was then incubated in a thermal cycler (BioRad, California, USA) according to the following parameters: priming at 25^0^C for 5 minutes; followed by reverse transcription at 46^0^C for 20 minutes and reverse transcription inactivation at 95^0^C for 1 minute. The synthesized cDNA was stored at -20^0^C.

b. qPCR was performed using the iTaq™ Universal SYBR Green Supermix (BioRad, California, USA) according to manufacturer’s protocol. The primers used were custom designed considering the gRNA oligo cute sites for both the genes. The primers were designed to determine the deletion of human *MCL-1* or *BFL-1* anti-apoptotic gene. Glyceraldehyde 3-phosphate dehydrogenase (*GAPDH*) was used as the housekeeping gene. The parental cells served as control.

***MCL-1:***

Forward oligo: 5^’^ – GCG GTA ATC GGA CTC AAC CT – 3^’^

Reverse oligo: 5^’^ – GTA GCC AAA AGT CGC CCT CC – 3^’^

***BFL-1:***

Forward oligo: 5^’^ – GCT GGC TCA GGA CTA TCT GC – 3^’^

Reverse oligo: 5^’^ – TGG ACG TTT TGC TTG GAC CT – 3^’^

***GAPDH:***

Forward oligo: 5^’^ – GTC TCC TCT GAC TTC AAC AGC G – 3^’^

Reverse oligo: 5^’^ – ACC ACC CTG TTG CTG TAG CCA A – 3^’^

c. All qPCR runs were carried out in the BioRad CFX96 qPCR platform. The samples were run in three biological and technical replicates. The reaction was set-up as below:

5 μl iTaq™ Universal SYBRR Green Supermix (2X)

**X** μl Forward and reverse primers (500 nM each)

**X** μl cDNA Variable (~10 ng)

Top up Nuclease-free water up to 10 μl

The thermal cycling protocol is as listed below:

| **Steps** | **Temperature**  **(ºC)** | **Time** | **No of cycles** |
| --- | --- | --- | --- |
| Polymerase activation and DNA denaturation | 95 | 2 minutes | 1 |
| Denaturation  Annealing | 95  55 | 15 seconds  30 seconds | 40 |

After thermal cycling, the PCR machine (BioRad CFX96) performed melt curve analysis. The melt-curve analysis was performed according to the instrument default settings.

**Two-dimensional (2D) Cytotoxicity Assay conducted with either *MCL-1* or *BFL-1* knock-out cells**

The HK-1 parental cell line, HK-1 sg*MCL-1*#2 and the HK-1 sg*BFL-1*#1 knock-out cells generated were treated with either ABT-199 or A1331852 in two-fold increasing doses (0.25-32 µM). After drug treatment, plates were incubated for 3-4 days in a humidified incubator at 37^0^C with 5% CO₂ and the assay was terminated, when cells in the untreated wells reached 80% to 90% confluency. After assay termination, the medium was “flicked off” from the plates and the plates were placed in freezer bags and stored in a -20°C freezer overnight. Next day, 200 μl of SyBr Green I dye (Thermo Fisher Scientific, MA USA) diluted in 1X Triton-X hypotonic lysis buffer at 1:4000 ratios were added to the 96-well plates. Plates were wrapped in aluminum foil as the SYBR Green I dye is light-sensitive. The stained plates were stored at 4°C for 4 days. After 4 days, cell proliferation was quantified using the CLARIOstar^®^ plate reader (BMG Labtech, Ortenberg, Germany) at 495 nm excitation and 530 nm emission filters.

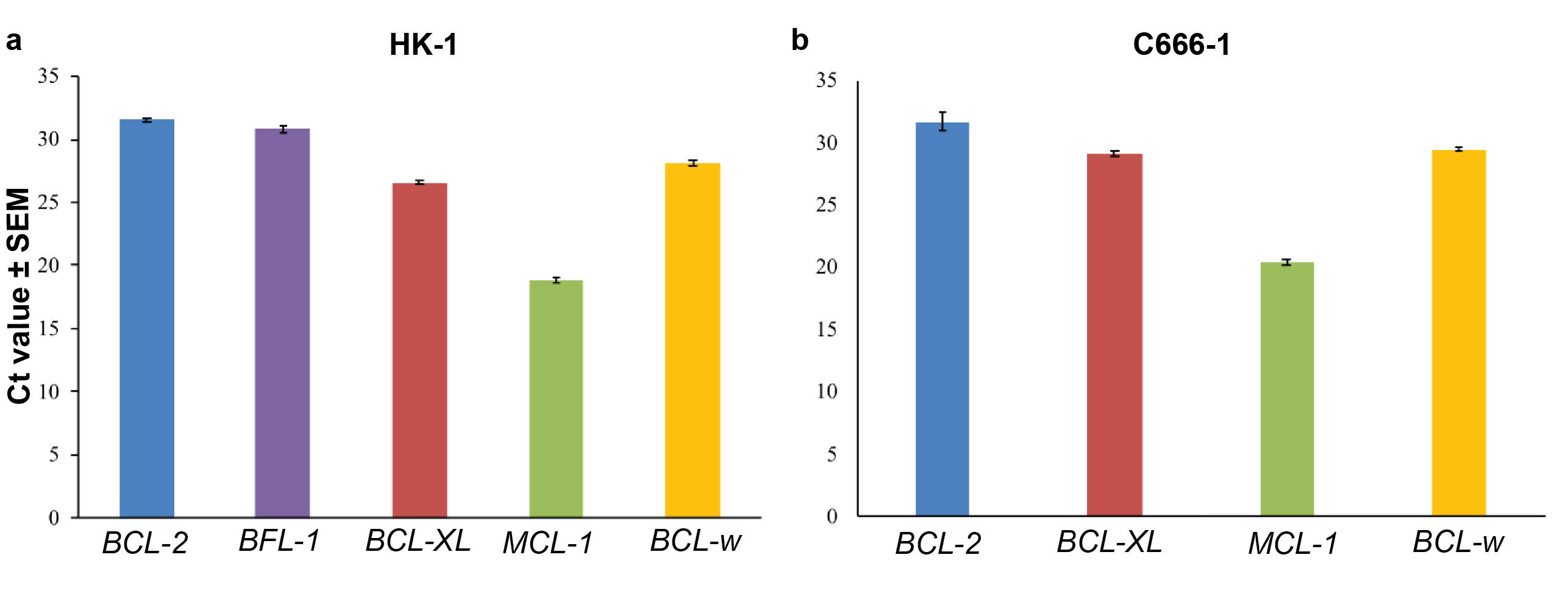

**S. Fig 3:** Expression levels of the anti-apoptotic genes in **(a)** HK-1 and **(b)** C666-1 NPC cell lines. Error bars show standard error of the mean (SEM).

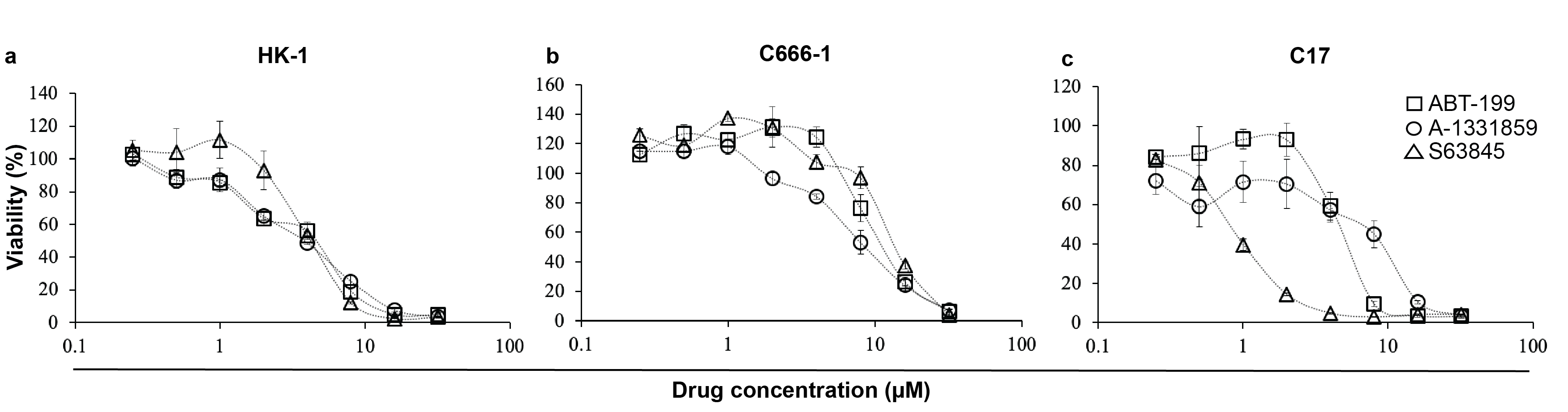

**S. Fig 4:** Treatment of **(a)** HK-1; **(b)** C666-1 and **(c)** C17 with single agent treatment of either ABT-199, A-1331852 or S63845 for 72 hours. Cell viability was assessed using the SyBr Green I assay. Points represent mean ± SEM of four experiments.

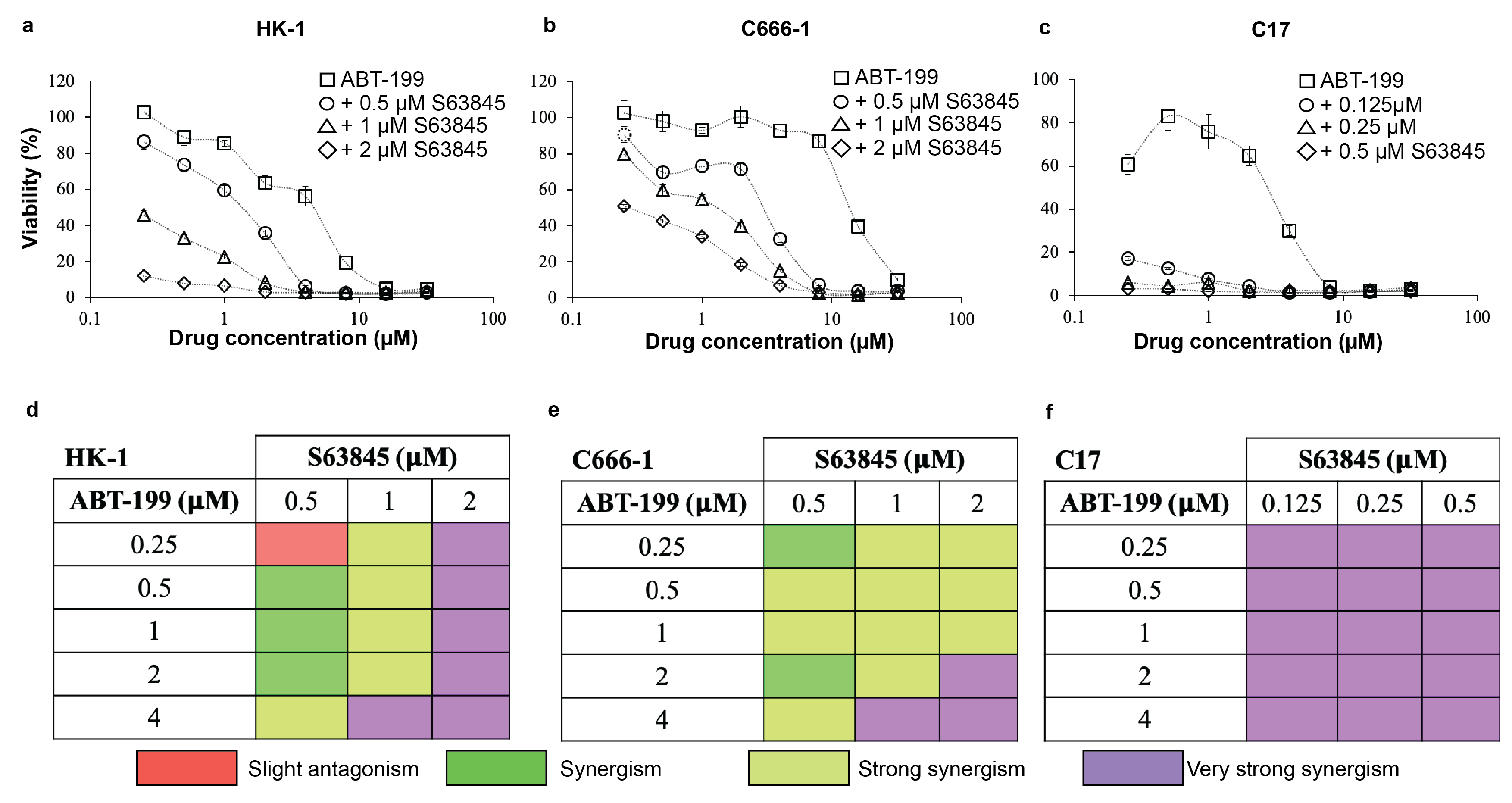

**S. Fig 5:** Treatment of **(a)** HK-1; **(b)** C666-1 and **(c)** C17 with single agent treatment of either ABT-199, A-1331852 or S63845 for 72 hours. Cell viability was assessed using the SyBr Green I assay. Points represent mean ± SEM of four experiments.

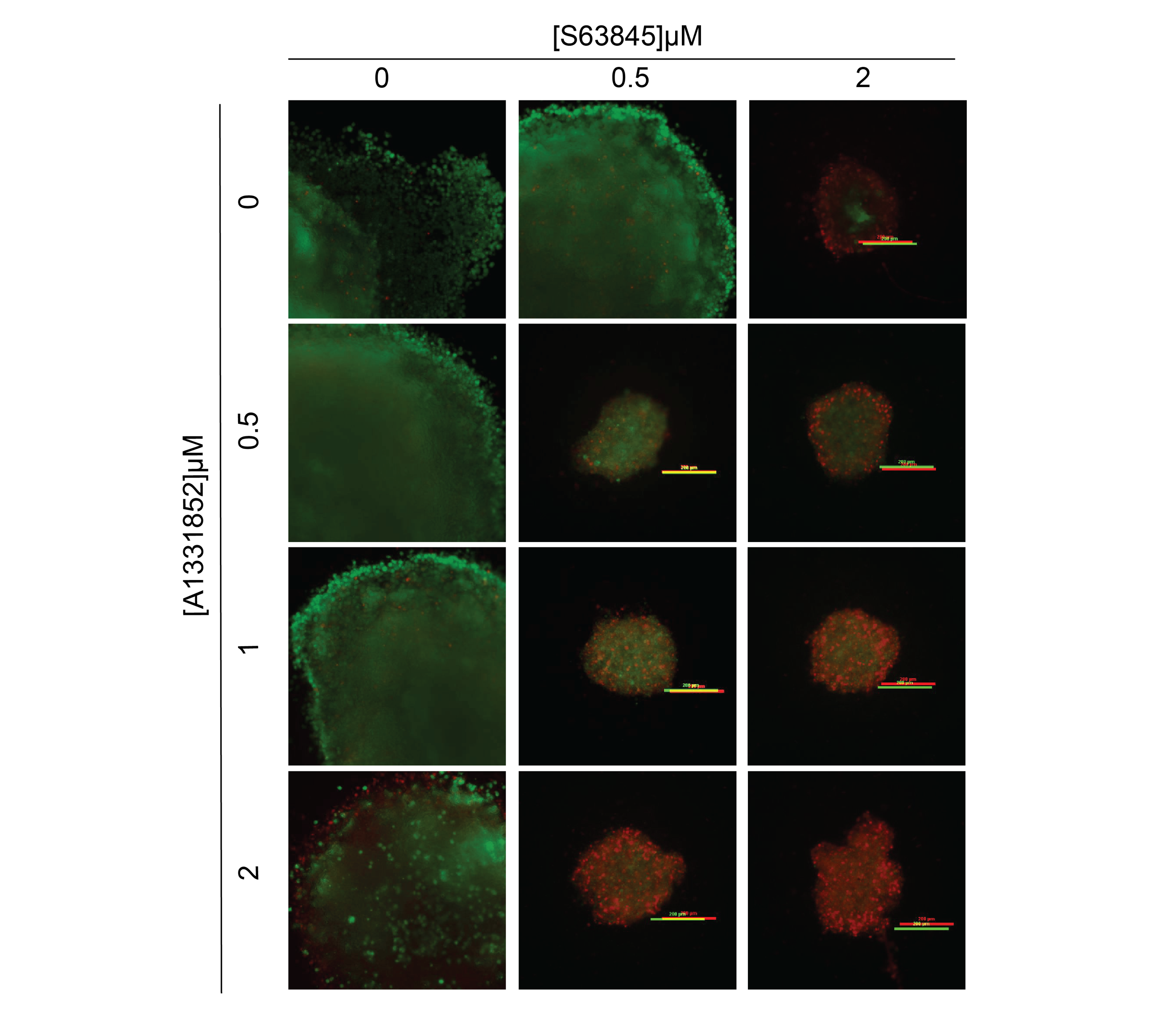

**S. Fig 6: The effect of combination of S63845 and A-1331852 on the growth and invasion of HK-1 spheroids over ten days.** The spheroids were treated with single agents S63845 and A-1331852 and combination of both over ten days at the indicated concentrations, n=2–3 spheroids per combination. Cell viability was determined using the live/dead assay (Viable cells: stained green by Calcein-AM; Dead cells: stained red by Ethidium-homodimer I). Images of viable cells and dead cells have been merged in this figure. Given that the growth of spheroids in the single agent panel were big, only part of the spheroid is shown for each drug concentration. Size bar: 200 μm.

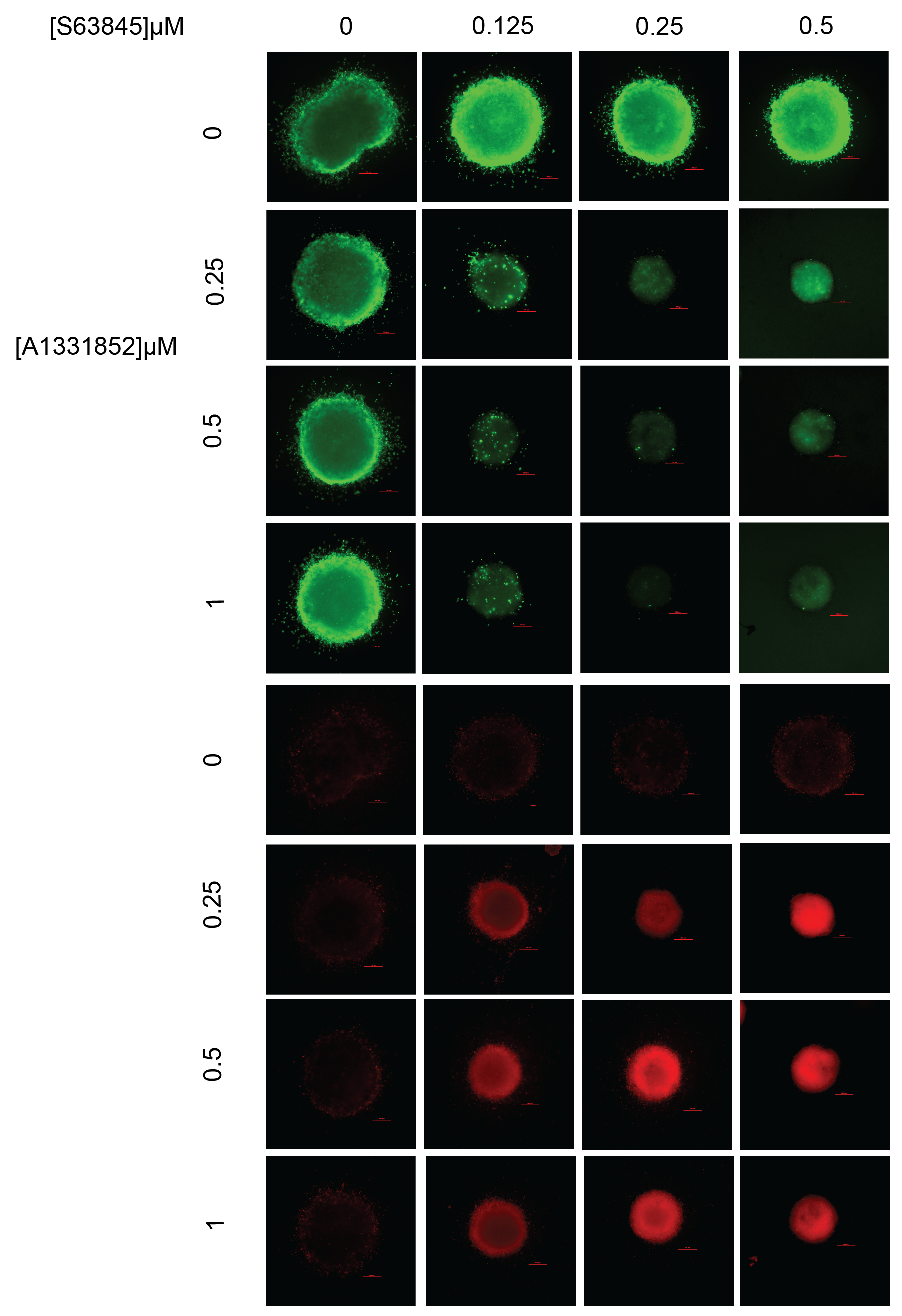

**S. Fig 7: The effect of combination of S63845 and A-1331852 on the growth and invasion of C17 spheroids over ten days.** The spheroids were treated with single agents S63845 and A-1331852 and combination of both over three days at the indicated concentrations, n=2–3 spheroids per combination. Cell viability was determined using the live/dead assay (Viable cells: stained green by Calcein-AM; Dead cells: stained red by Ethidium-homodimer I). Size bar: 200 μm.

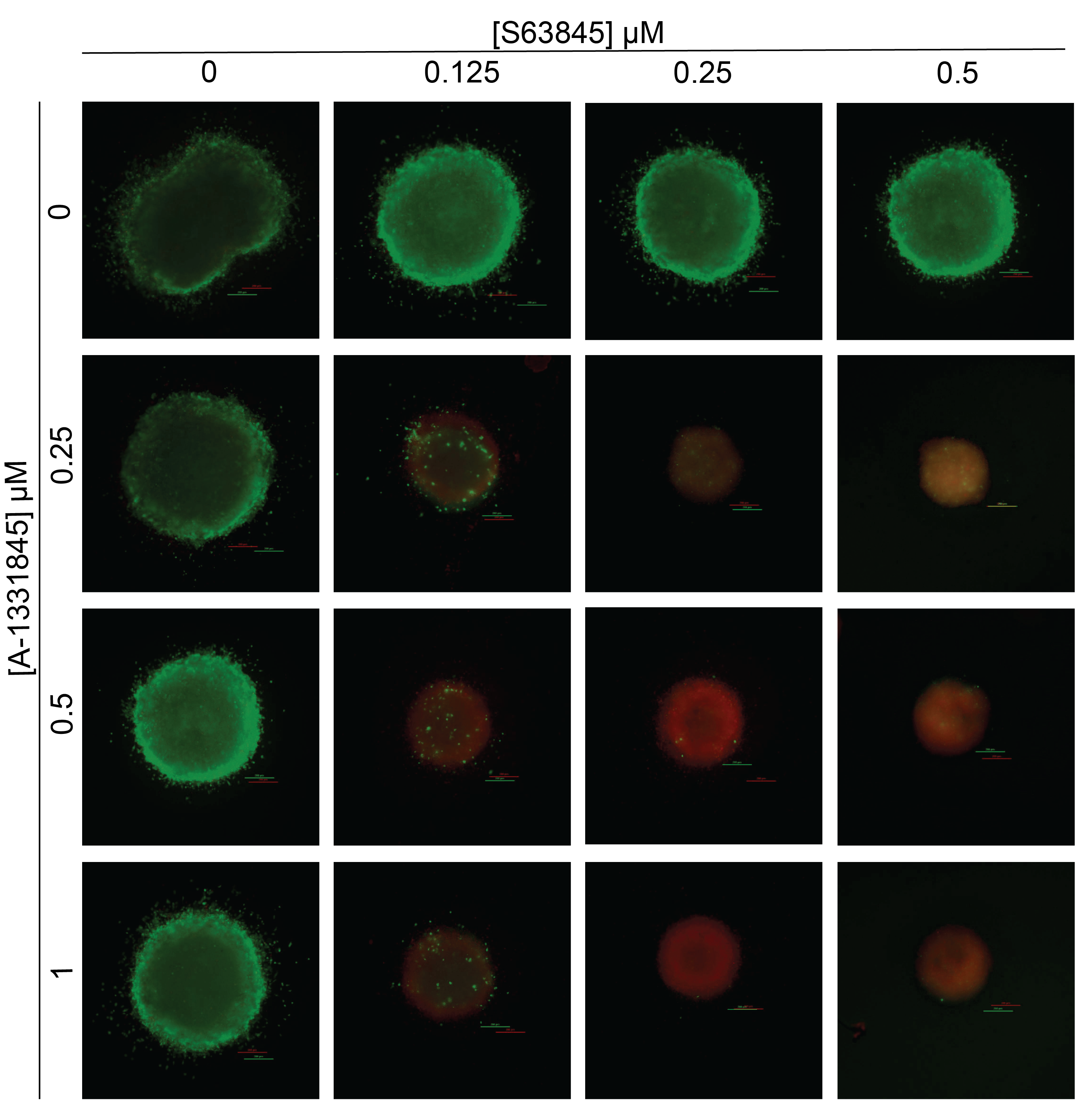

**S. Fig 8: The effect of combination of S63845 and A-1331852 on the growth and invasion of C17 spheroids over ten days.** The spheroids were treated with single agents S63845 and A-1331852 and combination of both over ten days at the indicated concentrations, n=2–3 spheroids per combination. Cell viability was determined using the live/dead assay (Viable cells: stained green by Calcein-AM; Dead cells: stained red by Ethidium-homodimer I). Images of viable cells and dead cells have been merged in this figure. Size bar: 200 μm.

**Supplementary table 1: Sensitization of NPC cell lines to ABT-199 by S63845 (fold sensitization).** The IC_50_ values are concentrations of drug 1 **(bold)** that kill 50% of the cells surviving the shown concentrations of *drug 2 (italics)*. Fold sensitization: IC_50_ drug 1/ IC_50_ drug 2.

| **Cell line** | *S63845 (µM)* | **ABT-199**  IC_50_ ± SEM (µM) | **Fold sensitization by S63845** |
| --- | --- | --- | --- |
| **HK-1** | 0 | 4.2 ± 0.65 | - |
|  | 0.5 | 1.3 ± 0.1 | 3.2 |
|  | 1 | 0.26 ± 0.005 | 16.1 |
|  | 2 | <0.25* | 16.8 |
| **C666-1** | 0 | 13.8 ± 0.63 | - |
|  | 0.5 | 2.7 ± 0.36 | 5.1 |
|  | 1 | 1.2 ± 0.25 | 11.6 |
|  | 2 | 0.38± 0.11 | 36.3 |
| **C17** | 0 | 2.6 ± 0.13 | - |
|  | 0.125 | <0.25 | 10.5 |
|  | 0.25 | <0.25* | 10.5 |
|  | 0.5 | <0.25* | 10.5 |

**Supplementary table 2:** Sensitivity of the HK-1 NPC cell line to either ABT-199 or A-1331852 following manipulation of *BFL-1*.

| **Drug** | **Cell Type** | **IC_50_ ± SEM (µM)** | **Fold sensitization** |
| --- | --- | --- | --- |
| ABT-199 | Parental HK-1 cell line | 5.28 ± 0.12 |  |
|  | HK-1 sg*BFL-1*#1 cells | 4.29 ± 0.36 | 1.2 |
| A-1331852 | Parental HK-1 cell line | 3.06 ± 0.23 |  |
|  | HK-1 sg*BFL-1*#1 cells | 0.67 ± 0.12 | 4.5 |

NOTE: Values were derived from at least 4 experiments. Sensitization was computed relative to the parent cell line, as shown.
